# Variability of Accessory Proteins Rules the SARS-CoV-2 Pathogenicity

**DOI:** 10.1101/2020.11.06.372227

**Authors:** Sk. Sarif Hassan, Pabitra Pal Choudhury, Vladimir N. Uversky, Guy W. Dayhoff, Alaa A. A. Aljabali, Bruce D. Uhal, Kenneth Lundstrom, Nima Rezaei, Murat Seyran, Damiano Pizzol, Parise Adadi, Amos Lal, Antonio Soares, Tarek Mohamed Abd El-Aziz, Ramesh Kandimalla, Murtaza Tambuwala, Gajendra Kumar Azad, Samendra P. Sherchan, Wagner Baetas-da-Cruz, Kazuo Takayama, Ángel Serrano-Aroca, Gaurav Chauhan, Giorgio Palu, Adam M. Brufsky

**Affiliations:** Department of Mathematics, Pingla Thana Mahavidyalaya, Maligram 721140, India; Applied Statistics Unit, Indian Statistical Institute, Kolkata 700108, West Bengal, India; Department of Molecular Medicine, Morsani College of Medicine, University of South Florida, Tampa, FL 33612, USA; Department of Chemistry, College of Art and Sciences, University of South Florida, Tampa, FL 33620, USA; Department of Pharmaceutics and Pharmaceutical Technology, Yarmouk University-Faculty of Pharmacy, Irbid 566, Jordan; Department of Physiology, Michigan State University, East Lansing, MI 48824, USA; PanTherapeutics, Rte de Lavaux 49, CH1095 Lutry, Switzerland; Research Center for Immunodeficiencies, Pediatrics Center of Excellence, Children’s Medical Center, Tehran University of Medical Sciences, Tehran, Iran & Network of Immunity in Infection, Malignancy and Autoimmunity (NIIMA), Universal Scientific Education and Research Network (USERN), Stockholm, Sweden; Doctoral studies in natural and technical sciences (SPL 44), University of Vienna, Austria; Italian Agency for Development Cooperation - Khartoum, Sudan Street 33, Al Amarat, Sudan; Department of Food Science, University of Otago, Dunedin 9054, New Zealand; Division of Pulmonary and Critical Care Medicine, Mayo Clinic, Rochester, Minnesota, USA; Department of Cellular and Integrative Physiology, University of Texas Health Science Center at San Antonio, 7703 Floyd Curl Dr, San Antonio, TX 78229-3900, USA; Department of Cellular and Integrative Physiology, University of Texas Health Science Center at San Antonio, 7703 Floyd Curl Dr, San Antonio, TX 78229-3900, USA & Zoology Department, Faculty of Science, Minia University, El-Minia 61519, Egypt; Applied Biology, CSIR-Indian Institute of Chemical Technology Uppal Road, Tarnaka, Hyderabad-500007, Telangana State, India & Department of Biochemistry, Kakatiya Medical College, Warangal, Telangana, India; School of Pharmacy and Pharmaceutical Science, Ulster University, Coleraine BT52 1SA, Northern Ireland, UK; Department of Zoology, Patna University, Patna-800005, Bihar, India; Department of Environmental Health Sciences, Tulane University, New Orleans, LA, 70112, USA; Translational Laboratory in Molecular Physiology, Centre for Experimental Surgery, College of Medicine, Federal University of Rio de Janeiro (UFRJ), Rio de Janeiro, Brazil; Center for iPS Cell Research and Application, Kyoto University, Japan; Biomaterials and Bioengineering Lab, Translational Research Centre San Alberto Magno, Catholic University of Valencia San Vicente Mártir, c/Guillem de Castro 94, 46001 Valencia, Spain; School of Engineering and Sciences, Tecnologico de Monterrey, Av. Eugenio Garza Sada 2501 Sur, 64849 Monterrey, Nuevo Léon, Mexico; Department of Molecular Medicine, University of Padova, Via Gabelli 63, 35121, Padova, Italy; University of Pittsburgh School of Medicine, Department of Medicine, Division of Hematology/Oncology, UPMC Hillman Cancer Center, Pittsburgh, PA, USA

**Keywords:** ORF3a, ORF6, ORF7a, ORF7b, ORF8, ORF10, Pathogenicity, SARS-CoV-2

## Abstract

The coronavirus disease 2019 (COVID-19) is caused by the Severe Acute Respiratory Syndrome Coronavirus-2 (SARS-CoV-2) which is pandemic with an estimated fatality rate less than 1% is ongoing. SARS-CoV-2 accessory proteins ORF3a, ORF6, ORF7a, ORF7b, ORF8, and ORF10 with putative functions to manipulate host immune mechanisms such as interferons, immune signaling receptor NLRP3 (NOD-, LRR-, and pyrin domain-containing 3) inflammasome, inflammatory cytokines such as interleukin 1*β* (IL-1*β*) are critical in COVID-19 pathology. Outspread variations of each of the six accessory proteins of all complete proteomes (available as of October 26, 2020, in the National Center for Biotechnology Information depository) of SARS-CoV-2, were observed across six continents. Across all continents, the decreasing order of percentage of unique variations in the accessory proteins was found to be ORF3a>ORF8>ORF7a>ORF6>ORF10>ORF7b. The highest and lowest unique variations of ORF3a were observed in South America and Oceania, respectively. This finding suggests that the wide variations of accessory proteins seem to govern the pathogenicity of SARS-CoV-2, and consequently, certain propositions and recommendations can be made in the public interest.

## 1. Introduction

SARS-CoV-2 (Severe Acute Respiratory Syndrome Coronavirus-2), the causative agent for The coronavirus disease 2019 (COVID-19), the pandemic is ongoing with the estimated fatality rate less than 1% [1]. However, the World Health Organization (WHO), Health Emergencies Programme, Executive Director, Dr Michael Ryan, in 2020 October indicated that 760 million might have been infected with SARS-CoV-2 which makes the hypothetical fatality rate as 0.14% with approximately one million lives taken. SARS-CoV-2 is a member of Betacoronavirus (lineage B) genus and Sarbecovirus subgenus was suggested to diverge from the lineage of Bat Coronavirus (BatCoV) Ratg13 in 1969 with the 95% highest posterior density interval of the years 1930 to 2000 [2]. Amongst previous Human Coronaviruses (HuCoVs) Severe acute respiratory syndrome-related coronavirus (SARS-CoV) is the closest to SARS-CoV-2 that caused an endemic from 2002 to 2004 [2, 3]. SARS-CoV had 8 open reading frame (ORF) 3a, 3b, 6, 7a, 7b, 8a, 8b, and 9b which were suggested to have more intrinsic and secondary other than having primary roles in the cell cycle and cellular entry [4, 5]. For instance, the ORFs are transcripted through the second phase of replication by (+) subgenomic messenger RNAs that were transcripted by the viral replication transcription complex negative-sense viral RNA coded in the initial stages of SARS-CoV-2 infection [6]. Thus, due to their intrinsic nature accessory proteins are not positive-selection sites such as extrinsic and primary functional Spike protein receptor-binding domain or protease cleavage sites [7]. Since, clinical SARS-CoV-2 isolates had a high-frequency non-synonymous mutation, D614G, in their S protein, which increased host cell entry via ACE2 and Transmembrane Protease Serine 2 (TMPRSS2) [8]. Therefore, due to the intrinsic nature and secondary order in the viral transcription, we can expect less selective pressure and mutations on accessory proteins to reach high-frequency in the population is less expected. Thus, despite the 19 to 89 years of estimated genomic divergence between SARS-CoV-2 and Ratg13, the sequence identity on accessory proteins ORF3, ORF6, ORF7a, ORF7b, ORF8, ORF10 had 98.5, 100, 97.5, 97.6, 95, and 100%, respectively are very high or identical which is indicating somehow the direct ancestor of SARS-CoV-2 had been exposed to almost no selective pressure to manipulate its intermediate host immunity for many years until the primary Human infection in Wuhan (Fig.1 to Fig.6) [2].

**Figure 1:**
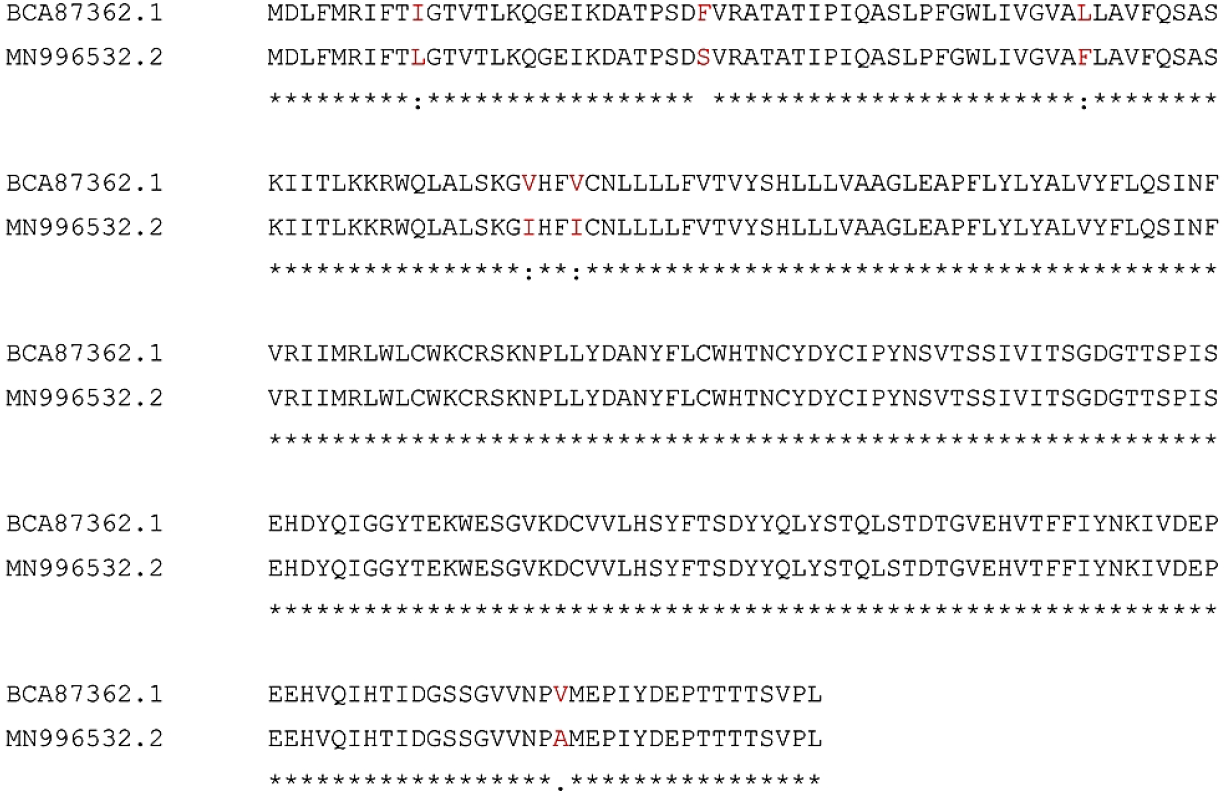
ClustalW alignment of SARS-CoV-2 and Ratg13 ORF3 proteins shows 98.5% sequence identity.

SARS-CoV-2 and SARS-CoV accessory proteins have differences such as putative protein ORF10 not present in SARS-CoV and the ORF3b and ORF9b is not present in SARS-CoV-2 [9, 10]. Very little is known about the functions of accessory proteins of SARS-CoV-2. The known essential features of the six accessory proteins are summarized below.

### ORF3a Protein

The ORF3a is the 275 amino acids long largest accessory protein among the accessory proteins coded by the SARS-CoV-2, has 72.4% sequence identity with SARS-CoV ORF3a protein and has 98.5% sequence identity with BatCoV Ratg13 ORF3a protein [11, 12] (Fig.1).

ORF3a is involved in virulence, infectivity, ion channel activity, morphogenesis, and virus release [13]. In SARS-CoV ORF3a was a multifunctional protein, co-localized with its protein binding regions with E, M, and S proteins in viral assembly formed homo-tetrameric complex as potassium-ion channel on host membrane [5]. In SARS-CoV-2, the ion-channel proteins (viroporins) function of ORF3a in addition to other proteins such as protein E, and ORF8a is critical in CoVs tissue inflammation [6]. Viroporins mediated lysosomal disruption and ion-redistribution activates innate immune signaling receptor NLRP3 (NOD-, LRR-, and pyrin domain-containing 3) inflammasome that leads to the expression of inflammatory cytokines such as interleukin 1), IL-6, and tumor necrosis factor (TNF), causing tissue inflammation during respiratory illness (IL[-61]. From another pathway, ORF3a with its protein binding domains interacts with TNF receptor-associated factor (TRAF3) protein, which leads ASC ubiquitination, and caspase 1 activation, and IL-1*β* maturation [14]. Additionally, ORF3a and ORF7a in combination with E, S, Nsp1 protein, and MAPK pathway proteins (MAPK8, MAPK14, and MAP3K7) trigger proinflammatory cytokine signaling transcription factors such as STAT1, STAT2, IRF9, and NFKB1 [6]. Another SARS-CoV-2 ORF3a protein interacts with heme oxygenase-1 (HMOX1) that has a role in heme catabolism and the anti-inflammatory system [6]. SARS-CoV-2 either triggers viral dissemination or suppresses continued viral replication of the apoptosis or programmed cell death [6]. In SARS-CoV ORF3a E and M protein, ion channel activity interferes with apoptotic pathways [11].

### ORF6 Protein

SARS-CoV-2 ORF6 is a 61 amino acid long membrane-associated interferon (IFN) antagonist protein suppresses the expression of co-transfected expression constructs and its subcellular localization to vesicular structures that has 68.9% sequence identity, with SARS-CoV ORF6 protein and has 100% sequence identity, with BatCoV Ratg13 ORF6 protein[5] (Fig.2).

**Figure 2:**
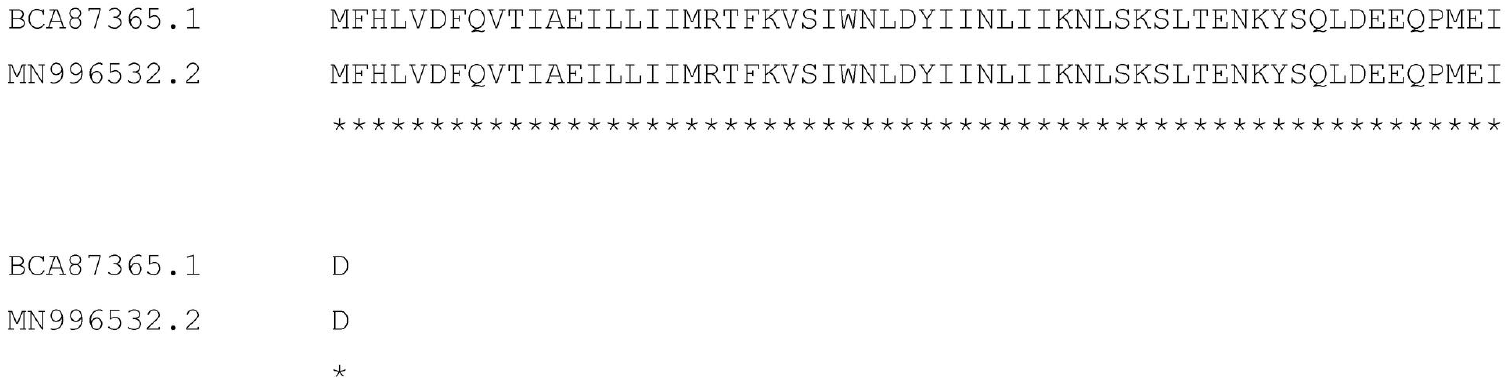
ClustalW alignment of SARS-CoV-2 (NCBI GenBank ID BCA87365.1) and Ratg13 (NCBI GenBank ID MN996532.2, translated 5 3 frame 1) ORF6 proteins shows 100% sequence identity, despite up to 89 years of genetic diversion.

ORF6 interacts with the karyopherin import complex that limits the movement of transcription factors STAT1 which down-regulates the IFN pathway [5]. In SARS-CoV ORF6 and ORF3a, in association with other proteins such as M, Nsp1 with Nsp3 inhibit IRF3 signalling and repress interferon expression and stimulate the degradation of IFNAR1 and STAT1 [6]. ORF6 interacts with the nsp8 protein coded by SARS-CoV-2 and it can increase infection during early infection at a low multiplicity with increase in RNA polymerase activity [15]. It is reported that ORF6 and ORF8 can inhibit the type-I interferon signaling pathway [15]. ORF6 protein with lysosomal targeting motif (YSEL) and diacidic motif (DDEE) induces intracellular membrane rearrangements resulting in a vesicular population and endosomal internalization of viral protein into the infected cells, increasing replication [16].

### ORF7a and ORF7b Proteins

ORF7a 121 aa coding type I transmembrane protein interacts with SARS-CoV-2 structural proteins M, E, and S, which are essential for viral assembly, and hence ORF7a is involved in the viral replication cycle and virion-associated ORF7a protein may function during early infection that has 85.2% Sequence identity with SARS-CoV ORF7a protein and has 97.5% sequence identity, with BatCoV Ratg13 ORF7a protein[5] (Fig.3).

**Figure 3:**
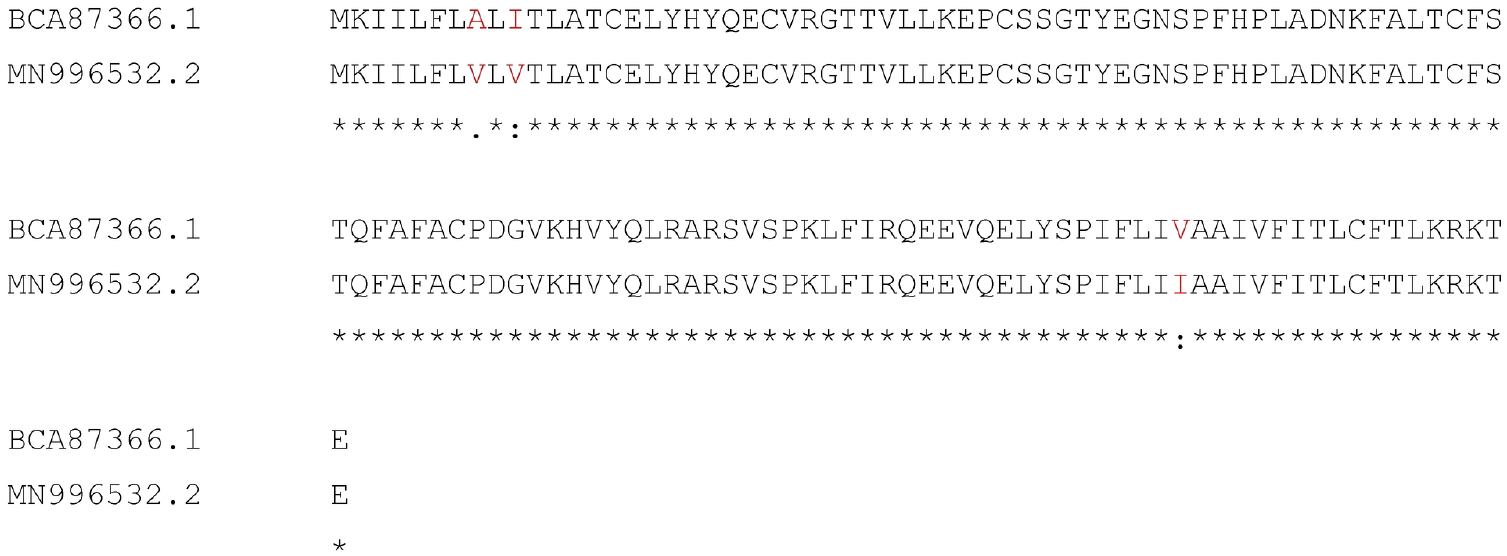
ClustalW alignment of SARS-CoV-2 (NCBI GenBank ID BCA87366.1) and Ratg13 (NCBI GenBank ID MN996532.2, translated 5’3’ frame 2) ORF7a proteins shows 97.5% sequence identity, despite up to 89 years of genetic diversion.

ORF7a interacts with SARS-CoV-2 structural proteins: membrane (M), envelope (E), and spike (S), which are essential for viral assembly, and hence ORF7a is involved in the viral replication cycle and virion-associated ORF7a protein may function during early infection [17, 18]. ORF7a leads the activation of pro-inflammatory cytokines and chemokines, such as IL-8 and RANTES [5]. SARS-CoV ORF7a in combination with E protein activate apoptosis by suppressing anti-apoptotic protein [6]. ORF7b is a 43 aa coding protein found in association with intracellular virus particles and also in purified virions inside the Golgi compartment that has an 85.4% sequence identity with SARS-CoV ORF7b protein and has 97.6% sequence identity, with BatCoV Ratg13 ORF7a protein [5] (Fig.4).

**Figure 4:**
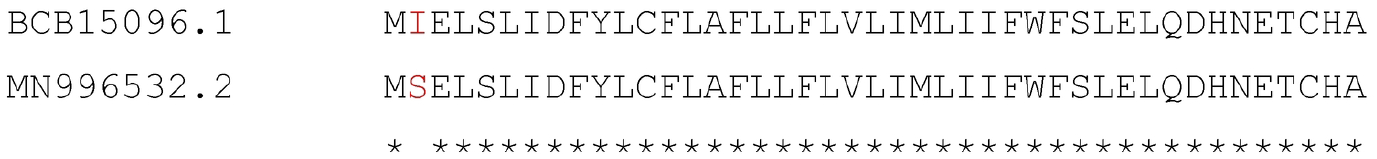
ClustalW alignment of SARS-CoV-2 (NCBI GenBank ID BCB15096.1) and Ratg13 (NCBI GenBank ID MN996532.2, translated 5’3’ frame 2) ORF7b proteins shows 97.6% sequence identity, despite up to 89 years of genetic diversion.

ORF7b is found in association with intracellular virus particles and also in purified virions. Till date, there is very little experimental evidence to support a role for ORF7a or ORF7b in the replication of SARS-CoV-2 [19].

### ORF8 Protein

ORF8 (121 aa long) is a unique accessory protein in SARS-CoV-2, and it stands out by being poorly conserved among other CoVs and accordingly showing structural changes suggested to be related to the ability of the virus to spread [20]. ORF8 is a unique accessory protein in SARS-CoV-2, which stands out by being poorly conserved among other CoVs and accordingly showing structural changes suggested to be related to the ability of the virus to spread [20]. ORF8 sequences of SARS-CoV-2 and Ratg13 share 95% amino acid identity (Fig.5).

**Figure 5:**
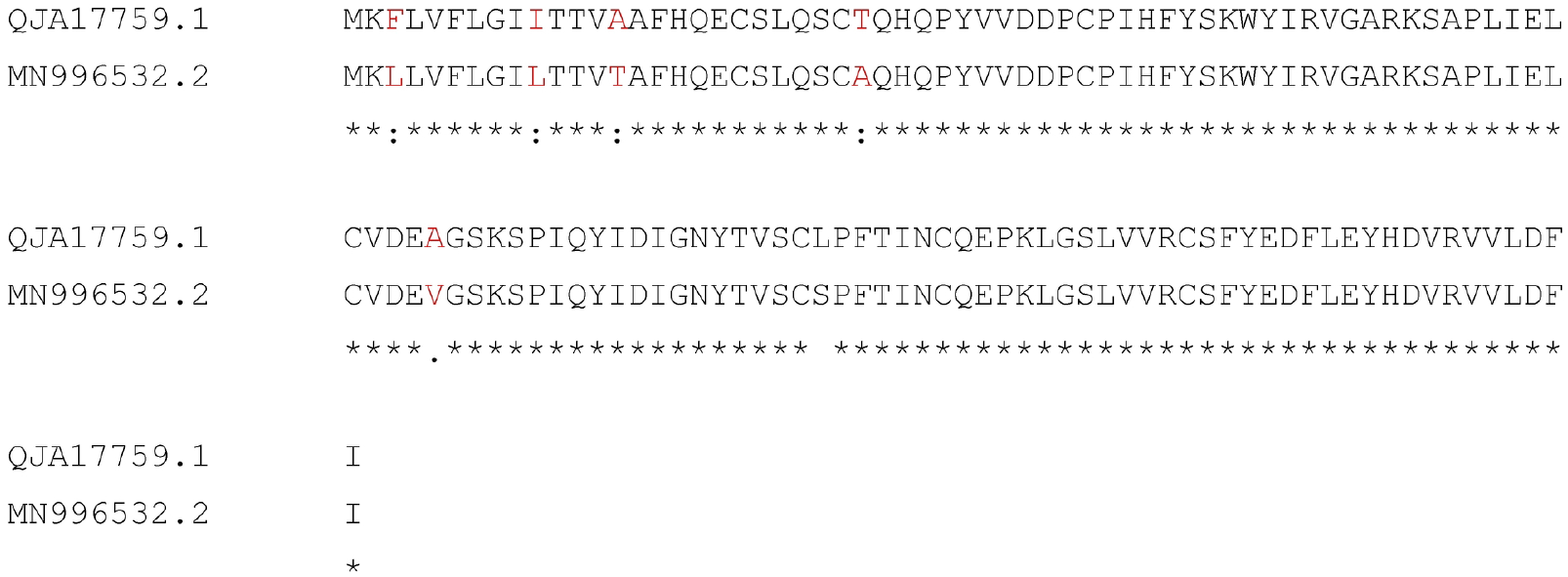
ClustalW alignment of SARS-CoV-2 (NCBI GenBank ID BCA87366.1) and Ratg13 (NCBI GenBank ID MN996532.2, translated 5’3’ frame 2) ORF8 proteins shows 95% sequence identity, despite up to 89 years of genetic diversion.

ORF8 of SARS-CoV-2 interacts with major histocompatibility complex (MHC) class-I molecules and down-regulates their surface expression significantly on various cell types [21]. It has been reported earlier that inhibition of ORF8 function could be a strategy to improve the special immune surveillance and accelerate the eradication of SARS-CoV-2 *in vivo* [22].

### ORF10 Protein

The accessory protein 38 aa coding protein ORF10 has been reported to be unique for SARS-CoV-2 containing eleven cytotoxic T lymphocyte (CTL) epitopes of nine amino acids in length each, across various human leukocyte antigen (HLA) subtypes [23, 24]. ORF10 negatively a ects the antiviral protein degradation process through its interaction with the Cul2 ubiquitin ligase complex [6]. SARS-CoV does not have ORF10 protein but SARS-COV-2 ORF10 and Ratg13 ORF10 has 97.3% sequence identity [25] (Fig.6).

**Figure 6:**
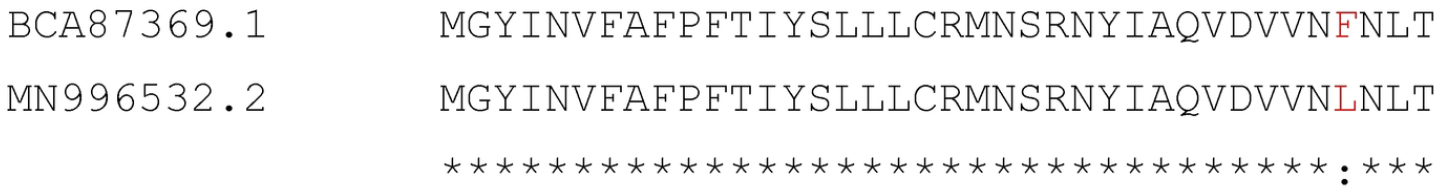
ClustalW alignment of SARS-CoV-2 (NCBI GenBank ID BCA87369.1) and Ratg13 (NCBI GenBank ID MN996532.2, translated 5’3’ frame 2) ORF10 proteins shows 97.3% sequence identity, despite up to 89 years of genetic diversion.

The objectives of the present study were to depict the unique variability of all accessory proteins and their possible contributions to virus pathogenicity.

## 2. Data acquisition

Sequences for all the accessory proteins ORF3a, ORF6, ORF7a, ORF7b, ORF8, and ORF10 were downloaded (on October, 20, 2020) from the complete SARS-CoV-2 proteomes on the National Center for Biotechnology Information (NCBI) database (http://www.ncbi.nlm.nih.gov/) (Table 1).

**Table 1:**
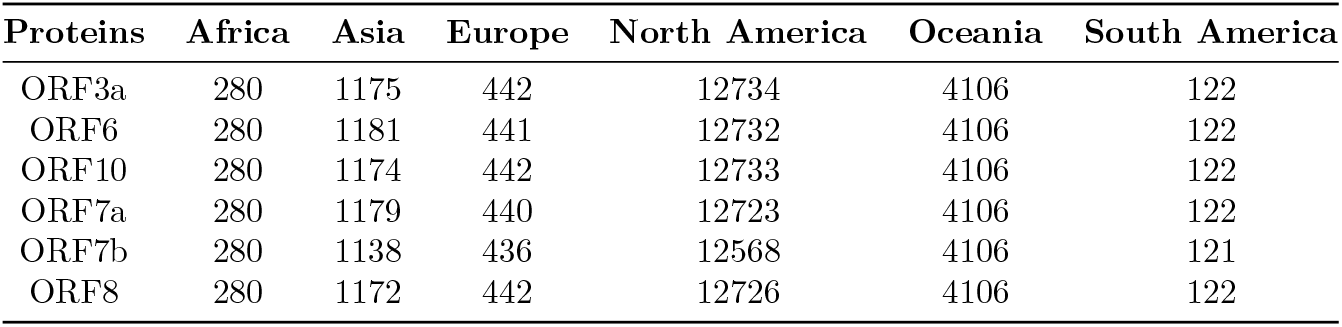
Total number of six accessory proteins of complete SARS-CoV-2 proteomes

| Proteins | Africa | Asia | Europe | North America | Oceania | South America |
| --- | --- | --- | --- | --- | --- | --- |
| ORF3a | 280 | 1175 | 442 | 12734 | 4106 | 122 |
| ORF6 | 280 | 1181 | 441 | 12732 | 4106 | 122 |
| ORF10 | 280 | 1174 | 442 | 12733 | 4106 | 122 |
| ORF7a | 280 | 1179 | 440 | 12723 | 4106 | 122 |
| ORF7b | 280 | 1138 | 436 | 12568 | 4106 | 121 |
| ORF8 | 280 | 1172 | 442 | 12726 | 4106 | 122 |

Note that all partial accessory proteins and sequences with ambiguous amino acids were excluded from the present study. Furthermore, the unique accessory protein sequences were extracted for each continent. The unique protein accessions were renamed for each accessory protein as S1, S2, … etc., as shown in the Supplementary Tables (7-13). There were 510, 72, 158, 37, 190, and 44 unique accessory proteins ORF3a, ORF6, ORF7a, ORF7b, ORF8, and ORF10, respectively, available. For each continent, ranges and names of sequences are presented in Table 2.

**Table 2:**
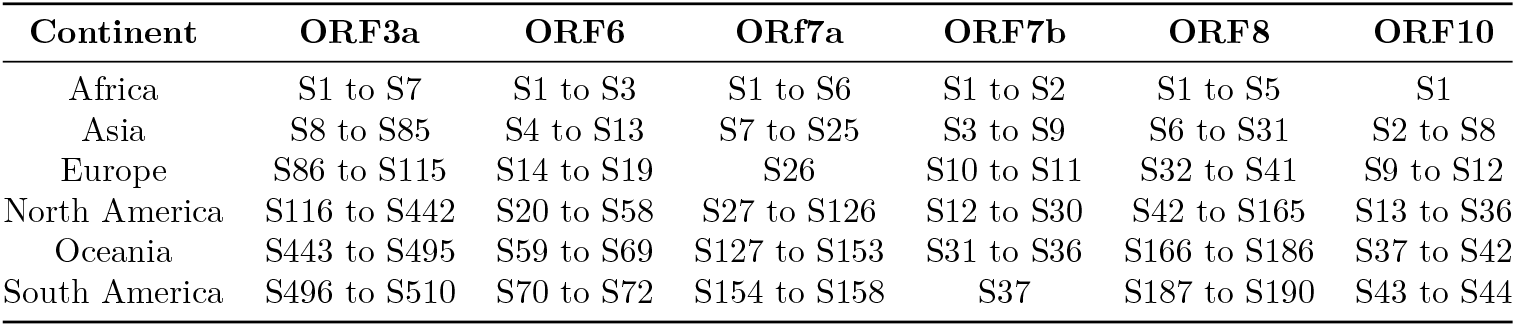
Ranges and naming of unique sequences (continent-wise) for each accessory protein of SARS-CoV-2

### 2.1. Evaluating the per-residue predisposition of SARS-CoV-2 accessory proteins and their natural variants for intrinsic disorder

Per-residue disorder distribution within the amino acid sequences of SARS-CoV-2 accessory proteins ORF3a, ORF6, ORF7a, ORF7b, ORF8 and ORF10 and their natural variants was evaluated by PONDR® VSL2, which is one of the more accurate standalone disorder predictors [26, 27, 28, 29]. The per-residue disorder predisposition scores are on a scale from 0 to 1, where values of 0 indicate fully ordered residues, and values of 1 indicate fully disordered residues. Values above the threshold of 0.5 are considered disordered residues, whereas residues with disorder scores between 0.25 and 0.5 are considered highly flexible, and residues with disorder scores between 0.1 and 0.25 are taken as moderately flexible.

## 3. Results

For every continent, the total number of accessory proteins and the total number of unique sequences with respective percentages are presented in Fig.7. In summary for all six continents, the total number of unique accessory proteins ORF3a, ORF6, ORF7a, ORF7b, ORF8, and ORF10 sequences are 419, 55, 122, 26, 147, and 32, respectively (Supplementary Fig.11). Furthermore, the percentage of unique sequences on each continent among all available accessory proteins are also enumerated (Fig.7).

**Figure 7:**
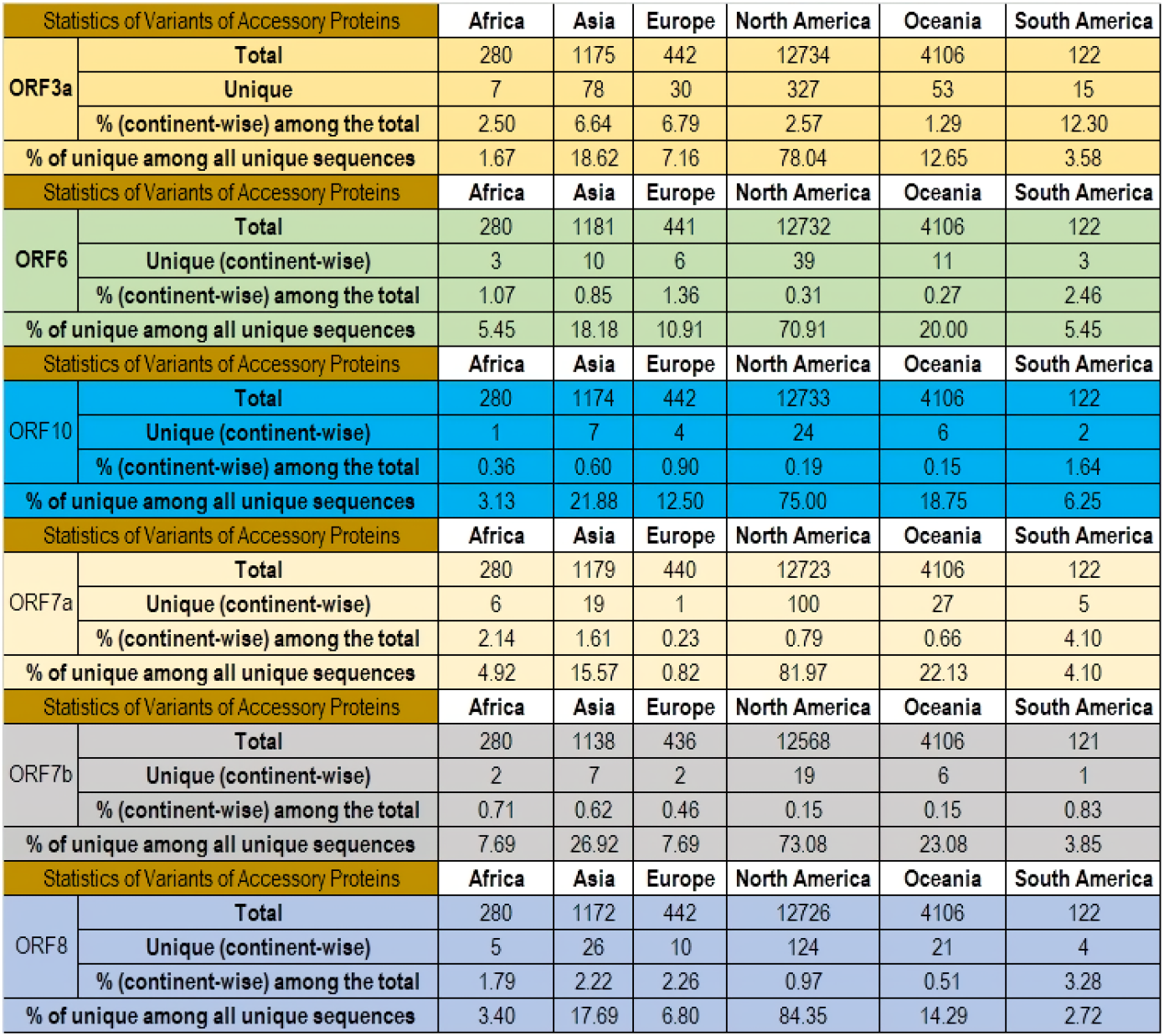
Number of unique accessory proteins across six continents

The percentage of each accessory protein across the six continents are presented as bar diagrams in Fig.8.

**Figure 8:**
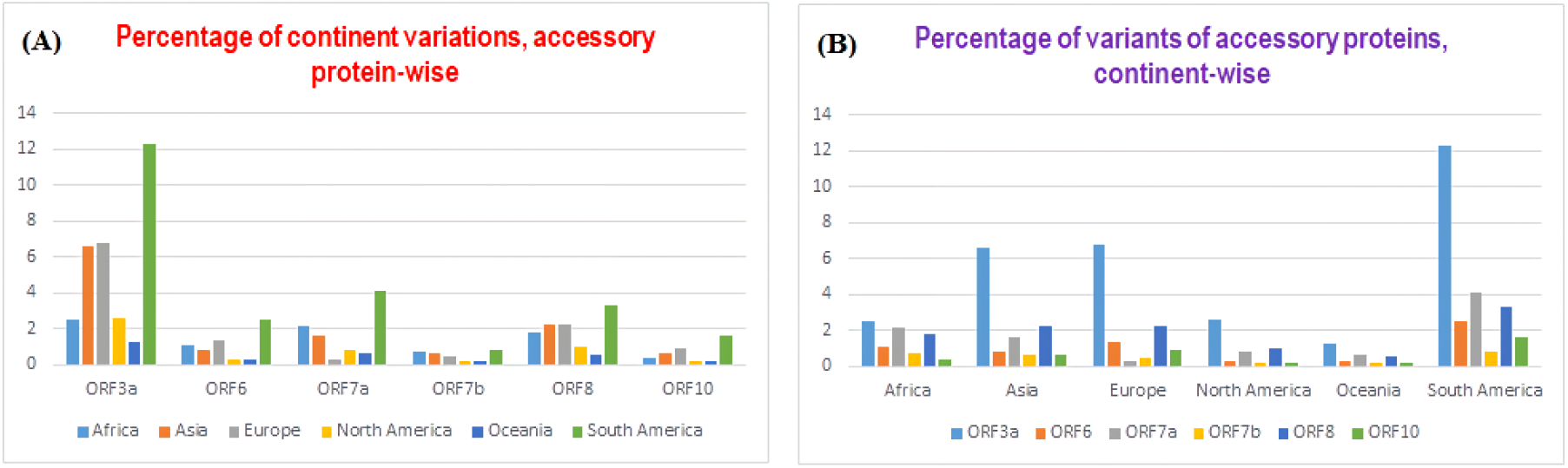
Bar representations of percentages of continental variations (A), and the percentage of unique accessory proteins (B).

From the Fig.8, the following observations were drawn:

Across all continents, the decreasing order of percentage of unique variations in the accessory proteins was observed as follows ORF3a>ORF8>ORF7a>ORF6>ORF10>ORF7b. The highest and lowest unique variations of ORF3a were observed in South America and Oceania, respectively. In addition, the highest percentage (statistically significant) of unique variations in each accessory protein was observed in South America. The lowest percentage of unique variations among ORF3a, ORF6, ORF7b, and ORF8 was observed in Oceania. It is worth noticing that the least number of unique variations in ORF7b and ORF7a was seen in North America and Europe, respectively. It is further noted that in Europe, the lowest variations among all accessory proteins was found in ORF7a. The smallest percentage of unique ORF10 variations was found in Oceania. With regards to the total unique variations across all accessory proteins of SARS-CoV-2, the decreasing order would be in South America>Asia>Europe>Africa>North America>Oceania. ORF3a possessed the highest amount (significantly) of unique variations across all the six continents while ORF10 showed the lowest variations in Africa, Asia, and Oceania. The lowest unique variations of ORF7b were observed in North America and South America. In addition, the percentage of unique accessory proteins among all unique sequences obtained across the six continents are represented as bar diagrams in Fig.9.

**Figure 9:**
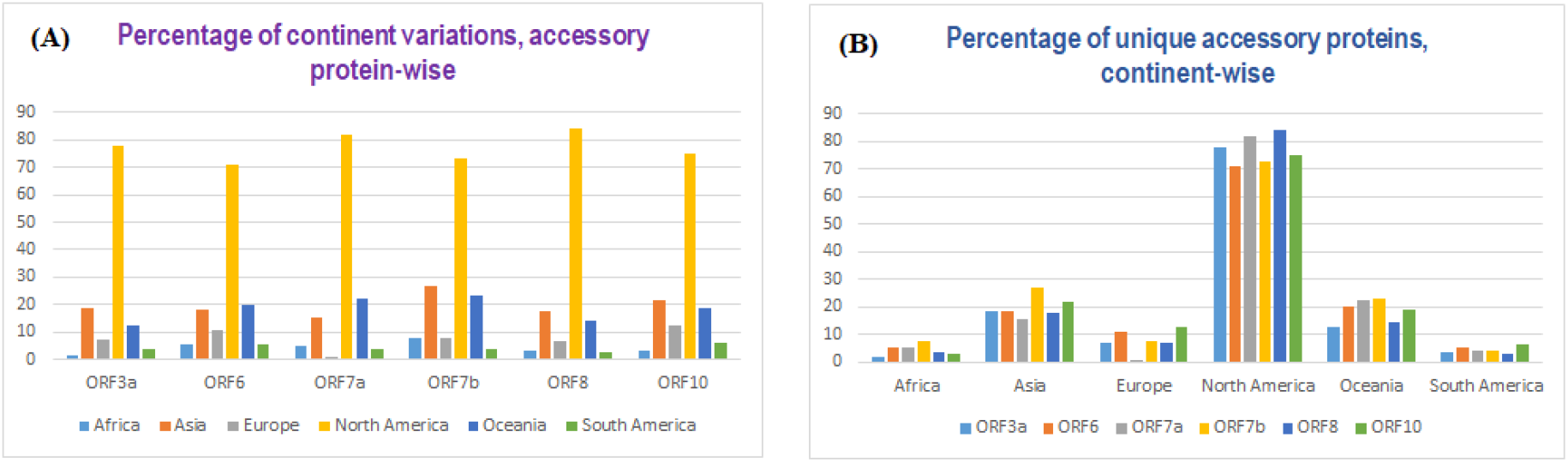
Quantitative information of the accessory proteins

The percentages of each unique variations among all unique variants of all accessory proteins across all six continents can be concluded from Fig.9 as follows:

Among all available unique variation of the six accessory proteins of SARS-CoV-2, North America and South America exhibited the highest and lowest percentage of variations of each accessory protein, respectively. The smallest number of unique variations of ORF3a, ORF6, and ORF10 were noticed in Africa. On the other hand, South America showed the lowest number of unique ORF6, ORF7a, ORF7b, and ORF8. With regards to ORF7b, the highest number of unique variations compared to the rest of the accessory proteins, were observed in Africa, Asia, and Oceania. Furthermore, the highest percentage (84.35%) and lowest (0.82%) of unique varieties of ORF8 and ORF7a (among all accessory proteins) was found in North America and Europe, respectively.

Following continent-wise, lists of identical sequences for each accessory protein were presented (Fig.10).

**Figure 10:**
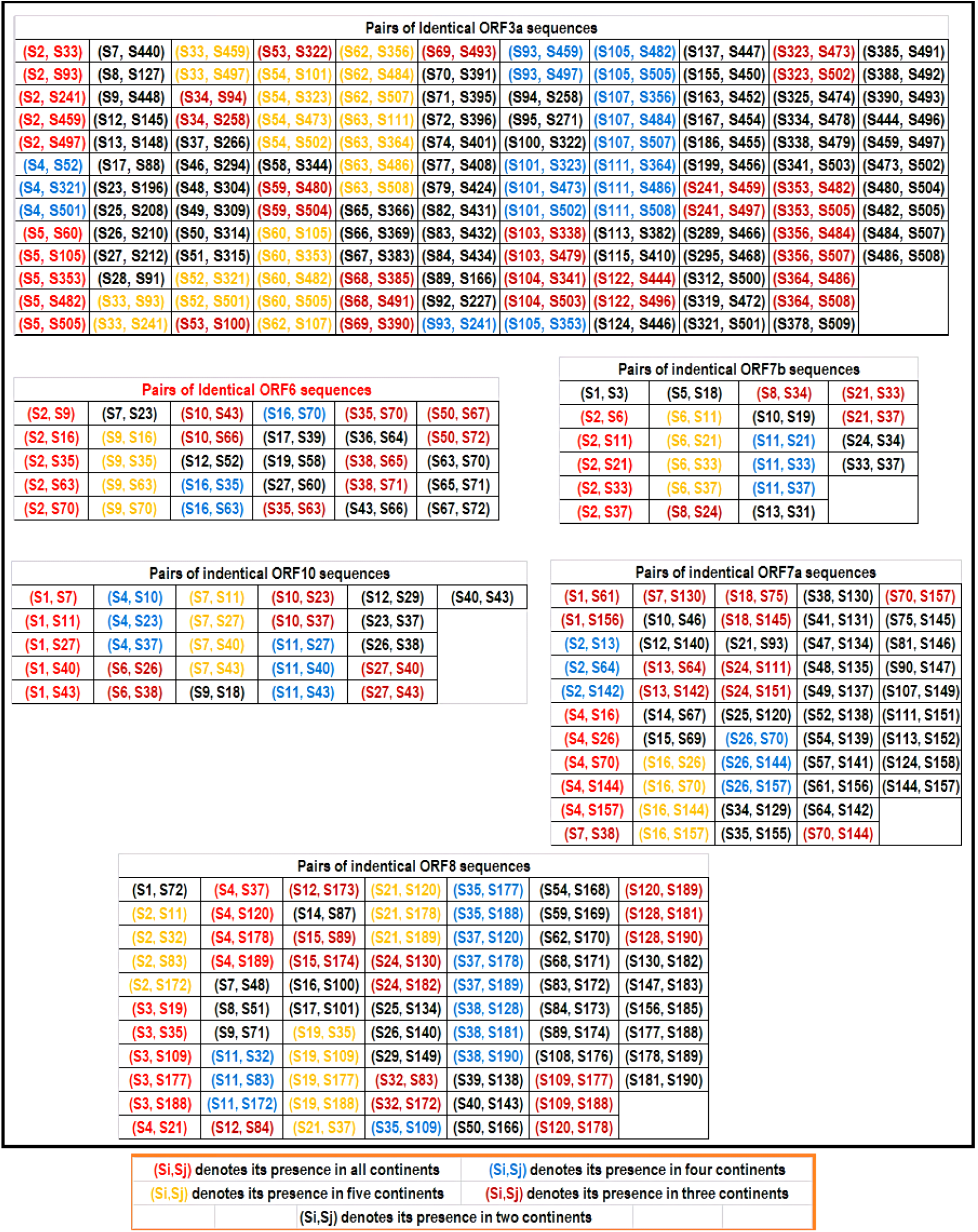
Identical pair of accessory protein sequences across all the continents.

The following observations were made for each accessory protein based on Fig.10:

### ORF3a

Note that, the mutations described below were determined based on the Wuhan ORF3a sequence (YP_009724391). There were only two ORF3a sequences (marked in red font), S2 (with reference to Africa, QOI60359) and S5 (with reference to Africa, QOI60335) which were present on all six continents. Note that the S2 (Africa-ORF3a) was identical with ORF3a (YP_009724391) from Wuhan, China. The other sequence S5 is different from ORF3a (YP_009724391) by one missense mutation Q57H, which was a strain determining mutation [30]. It is found that the ORF3a sequence S54 (Asia: QKK14624) possesses the single T175I mutation and is present on all continents except in Africa. The ORF3a sequences S62 (Asia: QMJ01306) and S63 (Asia: QJQ04482) possessed a single mutation each G251V and G196V, respectively with respect to Wuhan ORF3a (YP_009724391). These two sequences were present in Asia, Europe, North America, Oceania, and South America. The ORF3a sequence S4 (Africa: QLQ87565) has the single S171L mutation found on four continents excluding Europe and Oceania. Besides, two mutations Q57H and D155Y in sequence S34 (Asia) were present only on three continents, Asia, Europe, and North America. Sequence S53 (Asia) with the G172C mutation has been found in Asia, Europe and North

America only. The deletion mutation V255 occurred in S59 (Asia), which was found in Asia, Oceania, and South America. S68 (Asia) and S69 (Asia) possessed two mutations, H93Y and K67N, respectively. These two ORF3a variants have been detected only on three continents, Asia, North America, and Oceania. The ORF3a sequence S103 containing the single T229I mutation is present only on three continents, Europe, North America, and Oceania. Another sequence, S104, with the P240L mutation has been noticed only in Europe, North America, and South America. The V13L mutation was found in the S122 (ORF3a, North America) and is present on three continents, Oceania, North America, and South America. Further, there were 57 unique ORF3a variants detected only on two continents as listed in Table 3:

**Table 3:** List of ORF3a sequences and their presence on two continents only

| Sequence | Mutation(s) | Present in the continent(s) | Sequence | Mutation(s) | Present in the continent(s) |
| --- | --- | --- | --- | --- | --- |
| S7 | D2G | Asia and North America | S37 | Q57H, A103S | Asia and North America |
| S8 | L15F, Q57H | Asia and North America | S46 | L108F | Asia and North America |
| S9 | T32I | Asia and Oceania | S48 | W131C | Asia and North America |
| S12 | S40L, Q57H | Asia and North America | S49 | L140F | Asia and North America |
| S13 | L41F | Asia and North America | S50 | W149L | Asia and North America |
| S17 | V48F | Asia and Europe | S51 | T151I | Asia and North America |
| S23 | Q57H, W131C | Asia and North America | S58 | DEL(V255), N257D | Asia and North America |
| S25 | Q57H, S166L | Asia and North America | S65 | G172V | Asia and North America |
| S26 | Q57H, S171L | Asia and North America | S66 | D155Y | Asia and North America |
| S27 | Q57H, T175I | Asia and North America | S67 | A99V | Asia and North America |
| S28 | Q57H, S216P | Asia and Europe | S70 | K66N | Asia and North America |
| Sequence | Mutation(s) | Present in Continent(s) | Sequence | Mutation(s) | Present in Continent(s) |
| S71 | A54S, Q57H | Asia and North America | S167 | V55G | North America and Oceania |
| S72 | A54S | Asia and North America | S186 | Q57H, L101F | North America and Oceania |
| S74 | G49V | Asia and North America | S199 | Q57H, L140F | North America and Oceania |
| S77 | I35T, Q57H | Asia and North America | S289 | G100C | North America and Oceania |
| S79 | D22Y | Asia and North America | S295 | V112F | North America and Oceania |
| S82 | G18V, Q57H | Asia and North America | S312 | L147F | North America and South America |
| S83 | G18V | Asia and North America | S319 | S166L | North America and Oceania |
| S84 | K16N, Q57H | Asia and North America | S321 | S171L | North America and South America |
| S89 | V55F | Europe and North America | S325 | S177I | North America and Oceania |
| S92 | Q57H, V237F | Europe and North America | S334 | T223I | North America and Oceania |
| S94 | Q57H, D155Y | Europe and North America | S338 | T229I | North America and Oceania |
| S95 | Q57H, A99V | Europe and North America | S341 | P240L | North America and South America |
| S100 | G172C | Europe and North America | S378 | A110S | North America and South America |
| S113 | A39S | Europe and North America | S385 | H93Y | North America and Oceania |
| S115 | A33S, Q57H | Europe and North America | S388 | H78Y | North America and Oceania |
| S137 | S26L | North America and Oceania | S390 | K67N | North America and Oceania |
| S155 | L46F | North America and Oceania | S444 | V13L | Oceania and South America |
| S163 | L53F | North America and Oceania |  |  |  |

### ORF6

Note that the mutations described below were determined based on the Wuhan ORF6 sequence (YP_009724394). The sequence S2 (ORF6, Africa) was identical with YP_009724394 (China, Wuhan) ORF6 and this sequence was present on all six continents whereas ORF6 sequence, S10 (ORF6, Asia) with only the D53Y mutation, was found only in Asia, North America, and Oceania. The ORF6 sequences S38 (ORF6, North America) and S50 (ORF6, North America) possess a single mutation each, D2L and I33T, respectively, which were found on three continents, North America, Oceania, and South America. The ORF6 unique variant S7 (ORF6, Asia) possesses the E13D mutation, which was found only in Asia and North America. The ORF6 sequence S12 (ORF6, Asia) possessed a set of deletions “FKVSIWNLD” (22-30 aa) and it appeared in Asia and North America only. Sequence S17 (ORF6, Europe) had the D61Y mutation, and it was found in Europe and North America. In addition, a single mutation H3Y occurred in S19 (ORF6, Europe), which was present in Europe and North America. The ORF6 sequence S27 (ORF6, North America) containing the W27L mutation was found in North America and Oceania only. Furthermore, sequence S36 (ORF6, North America) with the D61H mutation was present in North America and Oceania only.

### ORF7a

Mutations are based on the Wuhan ORF7a sequence (YP_009724395). The Wuhan ORF7a sequence YP_009724395 was found on all continents. V104F was found in the sequence S2 (ORF7a, Africa) in Africa, Asia, North America, and Oceania. The sequence S1 (ORF7a, Africa) had the P39L mutation, which was found in Africa, North America, and South America. S37F was found in sequence S7 (ORF7a, Asia) in Asia, North America, and Oceania. Sequence S18 (ORF7a, Asia) has the A105V mutation found across Asia, North America, and Oceania. G38V was found in S24 (ORF7a, Asia) in Asia, North America, and Oceania. Also, there were 21 unique ORF7a variants, present only on two continents. All mutations are listed in Table 4:

**Table 4:**
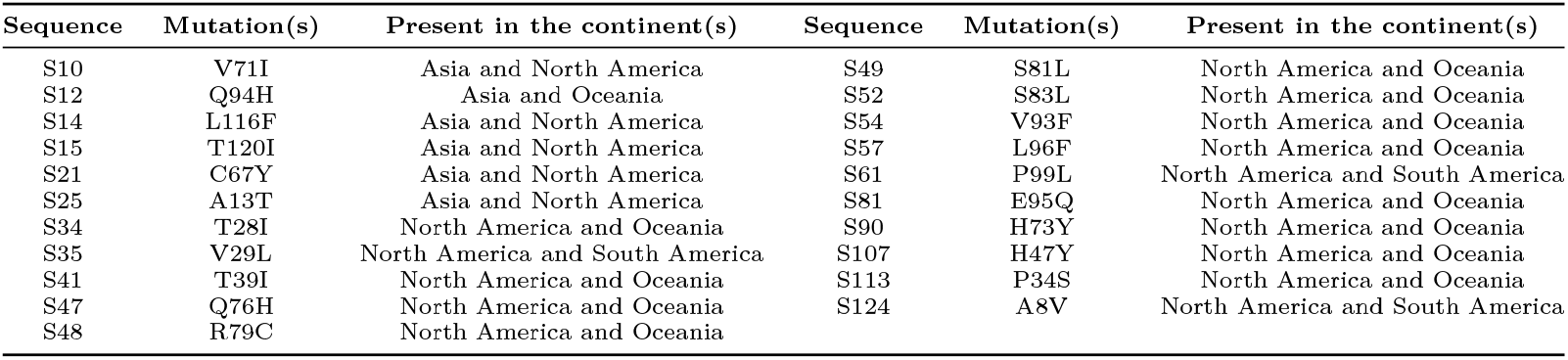
List of ORF7a sequences and their presence on two continents only

| Sequence | Mutation(s) | Present in the continent(s) | Sequence | Mutation(s) | Present in the continent(s) |
| --- | --- | --- | --- | --- | --- |
| S10 | V71I | Asia and North America | S49 | S81L | North America and Oceania |
| S12 | Q94H | Asia and Oceania | S52 | S83L | North America and Oceania |
| S14 | L116F | Asia and North America | S54 | V93F | North America and Oceania |
| S15 | T120I | Asia and North America | S57 | L96F | North America and Oceania |
| S21 | C67Y | Asia and North America | S61 | P99L | North America and South America |
| S25 | A13T | Asia and North America | S81 | E95Q | North America and Oceania |
| S34 | T28I | North America and Oceania | S90 | H73Y | North America and Oceania |
| S35 | V29L | North America and South America | S107 | H47Y | North America and Oceania |
| S41 | T39I | North America and Oceania | S113 | P34S | North America and Oceania |
| S47 | Q76H | North America and Oceania | S124 | A8V | North America and South America |
| S48 | R79C | North America and Oceania |  |  |  |

### ORF7b

Here all mutations are accounted based on the Wuhan ORF7b sequence (YP_009725318). The sequence S2 (ORF7b, Africa) (identical to Wuhan ORF7b (YP_009725318)) was found on all the six continents. It is found that only the C41F mutation was present in S8 (ORF7b, Asia) which appeared in Asia, North America, and Oceania. The sequence S1 (ORF7b, Africa) had the single mutation S5L and it was present in Africa and Asia. Sequence S5 (ORF7b, Asia) had the mutation S31L and this sequences was found on the two continents, Asia and North America only. L32F occurred in the sequence S10 (ORF7b, Europe) which was present on the continents Europe and North America. Furthermore, ORF7b Sequence S13 had the mutation L4F and this sequence was found on North America and Oceania.

### ORF8

Mutations described below are determined with reference to the Wuhan ORF7b sequence (YP_009724396). It is observed that, the Wuhan ORF8 YP_009724396 sequence was found on every continent. Also, there was another sequence which is also present in every continent, having the single mutation L84S. The V62L, a single mutation was observed in the sequence S2 (ORF8, Africa) which was found on all continents except South America, whereas ORF8 sequence S38 (Europe) possessed the single mutation A65S and the sequence was found in North America, Oceania, and South America. Further, V62L and L84S two mutations were observed in S12 (ORF8, Asia) and S12 appeared in Asia, North America, and Oceania. Sequence S15 (ORF8, Asia) got a mutation S67F and it was found in Asia, North America, and Oceania. ORF8 sequence S24 (Asia) possessed a single mutation A65V and the sequence was found in Asia, North America, and Oceania.

### ORF10

Here, mutations are based on the Wuhan ORF10 sequence (YP_009725255). The Wuhan ORF10 (YP_009725255) became identical with S1 (ORF10, Africa) and it was found on every continent. ORF10 sequence S6 (ORF10, Asia) had the mutation L37F and the sequence was present on North America, and Oceania only. The only mutation V30L was found in ORF10 sequence S10 (Europe) which appeared in Europe, North America, and Oceania. The sequence S9 (ORF10, Europe) had the mutation S23F and it was found in Europe and North America. Also, the mutation D31Y appeared in S12 (ORF10, Europe) which was found in Europe and North America only.

### 3.1. Featuring uniqueness of the accessory proteins

Here certain basic descriptive statistics (mean, variance, lower bound, upper bound, and range) were employed to describe the variability of the percentage of intrinsic protein disordered residues (IPD), molecular weight (MW), and isoelectric point (IP) of all the unique variants of all accessory proteins (Table 6). The zigzag behavior of the plots of IPD, MW, and IP depicts the wide variability of each accessory protein-variant (Supplementary Fig.12-Fig.15).

**Table 5:**
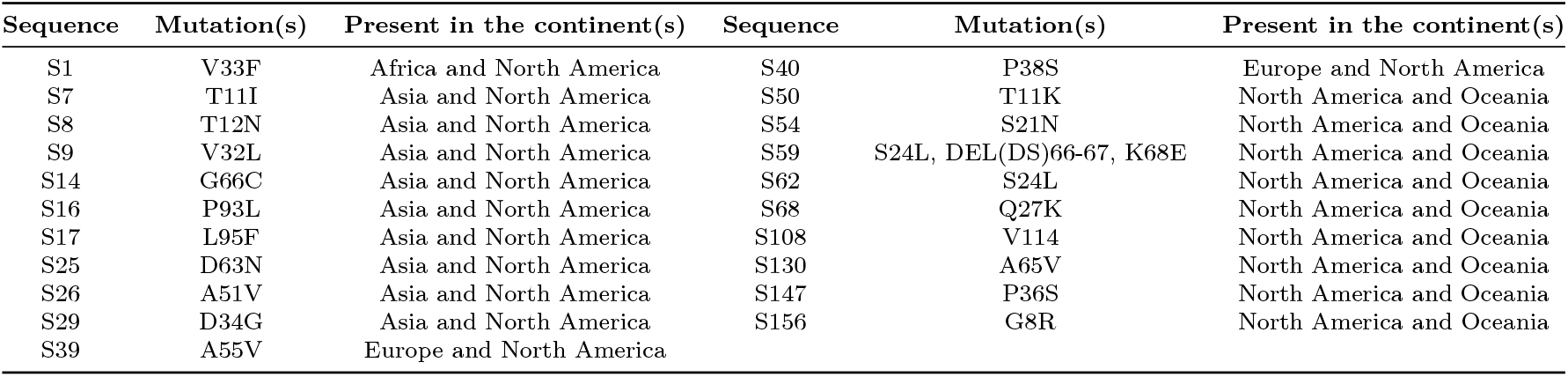
List of ORF8 sequences and their presence on two continents only

| Sequence | Mutation(s) | Present in the continent(s) | Sequence | Mutation(s) | Present in the continent(s) |
| --- | --- | --- | --- | --- | --- |
| S1 | V33F | Africa and North America | S40 | P38S | Europe and North America |
| S7 | T11I | Asia and North America | S50 | T11K | North America and Oceania |
| S8 | T12N | Asia and North America | S54 | S21N | North America and Oceania |
| S9 | V32L | Asia and North America | S59 | S24L, DEL(DS)66-67, K68E | North America and Oceania |
| S14 | G66C | Asia and North America | S62 | S24L | North America and Oceania |
| S16 | P93L | Asia and North America | S68 | Q27K | North America and Oceania |
| S17 | L95F | Asia and North America | S108 | V114 | North America and Oceania |
| S25 | D63N | Asia and North America | S130 | A65V | North America and Oceania |
| S26 | A51V | Asia and North America | S147 | P36S | North America and Oceania |
| S29 | D34G | Asia and North America | S156 | G8R | North America and Oceania |
| S39 | A55V | Europe and North America |  |  |  |

**Table 6:** Descriptive statistics of IPD, MW, and IP of unique accessory proteins of SARS-CoV-2

| IPD of unique accessory proteins of SARS-CoV-2 |  |  |  |  |  |
| --- | --- | --- | --- | --- | --- |
| *Accessory proteins* | *Mean* | *Variance* | *Lower bound* | *Upper bound* | *Range* |
| ORF3a | 4.756 | 0.2328 | 2.91 | 7.64 | 4.73 |
| ORF6 | 25.74 | 74.69 | 21.31 | 87.5 | 66.19 |
| ORF7a | 3.51 | 0.5716 | 2.48 | 7.29 | 4.81 |
| ORF7b | 44.663 | 10.527 | 37.21 | 51.16 | 13.95 |
| ORF8 | 9.125 | 1.285 | 5.6 | 13.45 | 7.85 |
| ORF10 | 18.67 | 5.0691 | 13.16 | 23.68 | 10.52 |
| MW of unique accessory proteins of SARS-CoV-2 |  |  |  |  |  |
| *Accessory proteins* | *Mean* | *Variance* | *Lower bound* | *Upper bound* | *Range* |
| ORF3a | 31123 | 17917.58 | 29187 | 31270 | 2083 |
| ORF6 | 7171.03 | 371714.6 | 2881.205 | 7542.84 | 4661.635 |
| ORF7a | 13673.4 | 150719.4 | 10874.515 | 14328.65 | 3454.135 |
| ORF7b | 5173.02 | 2651.26 | 5033.005 | 5224.22 | 191.215 |
| ORF8 | 13841.4 | 21411.43 | 12608.465 | 14431.55 | 1823.085 |
| ORF10 | 4446.53 | 1173.801 | 4389.085 | 4509.285 | 120.2 |
| IP of unique accessory proteins of SARS-CoV-2 |  |  |  |  |  |
| *Accessory proteins* | *Mean* | *Variance* | *Lower bound* | *Upper bound* | *Range* |
| ORF3a | 5.9127 | 0.0278 | 5.2349 | 6.5881 | 1.3532 |
| ORF6 | 4.4013 | 0.057 | 3.8436 | 5.7589 | 1.9153 |
| ORF7a | 8.0932 | 0.0434 | 6.7486 | 8.5946 | 1.846 |
| ORF7b | 3.9519 | 0.0063 | 3.6379 | 4.1442 | 0.5063 |
| ORF8 | 5.6368 | 0.1223 | 4.7442 | 6.8829 | 2.1387 |
| ORF10 | 8.2415 | 0.6857 | 6.0601 | 9.2043 | 3.1442 |

The following observations were made based on Table 6.

The amount total dispersions (based on range) of the percentage of IPD and MW of ORF6 variants turned out to be highest whereas the highest amount of total dispersion of IP was observed for ORF10. The smallest amount of total dispersions of the percentage of IPD, MW, and IP were found for ORF3a, ORF10, and ORF7b, respectively. The large value of range and variance of the MW of the unique ORF3a, ORF7a, ORF8, and ORF10 variants imply the wide variability of each set of ORF3a, ORF7a, ORF8, and ORF10 though range and variance of IPD and IP were not much widely spread. In case of unique variance of ORF6, the range and variance of MW and percentage of IPD were found to be large which implied the wide quantitative differences among the unique ORF6 variants. Furthermore, moderately high range and variance associated with the percentage of IPD and MW of ORF7a variants imply its moderate variability.

In line with the previously reported data, Fig.14 and Table 6 show that all SARS-CoV-2 accessory proteins contain different levels of intrinsic disorder. Furthermore, this analysis revealed that intrinsic disorder predispositions can vary significantly between the natural variants of each individual accessory protein. Importantly, the largest mutation-induced variability is observed within the disordered or flexible regions of these proteins (i.e., regions characterized by the predicted disorder scores exceeding the 0.5 threshold and regions with disorder scores between 0.25 and 0.5). This is an important observation suggesting that natural variability of SARS-CoV-2 accessory proteins is shaping their structural flexibility.

**Figure 11:**
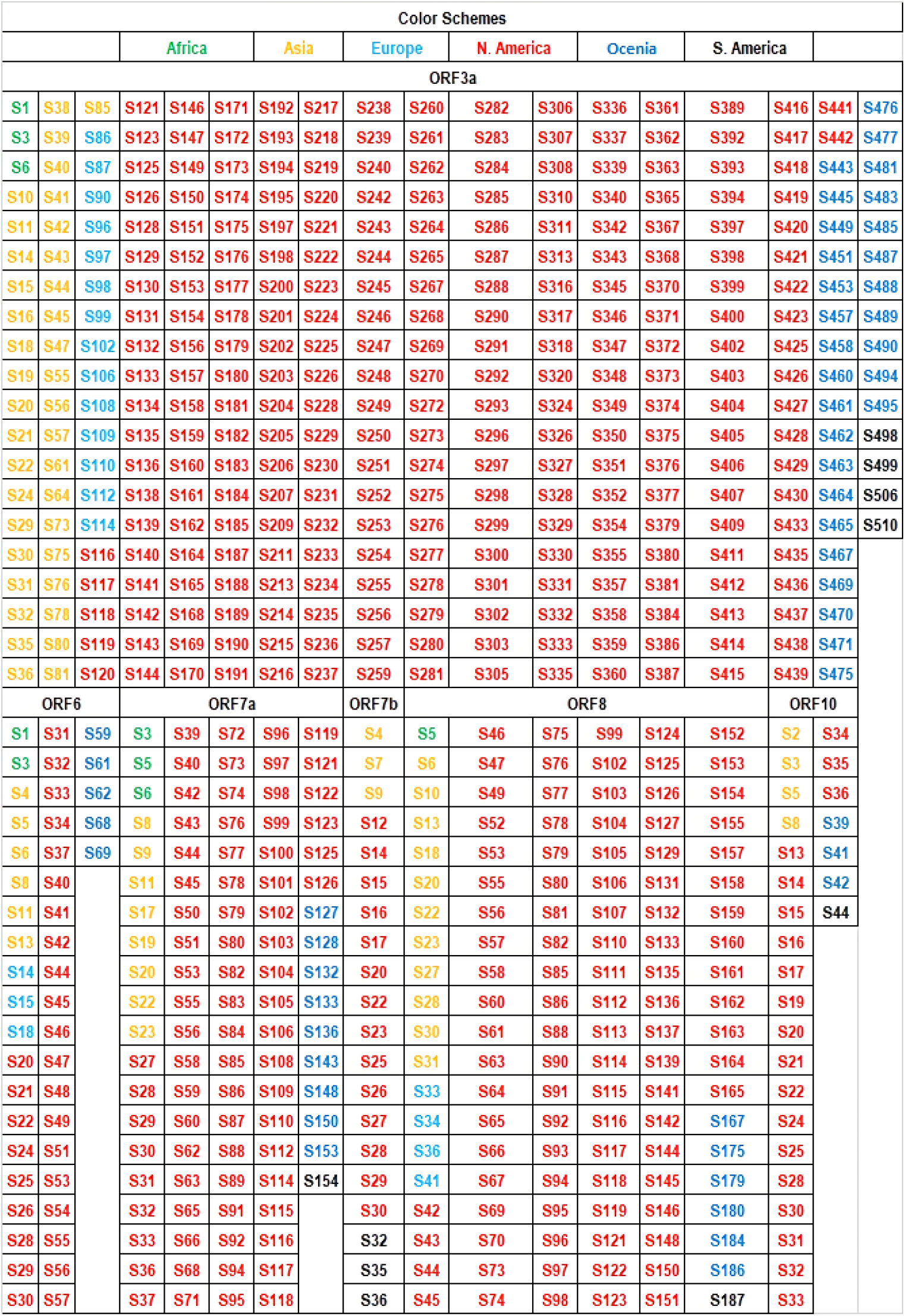
Unique variants of accessory protein sequences.

**Figure 12:**
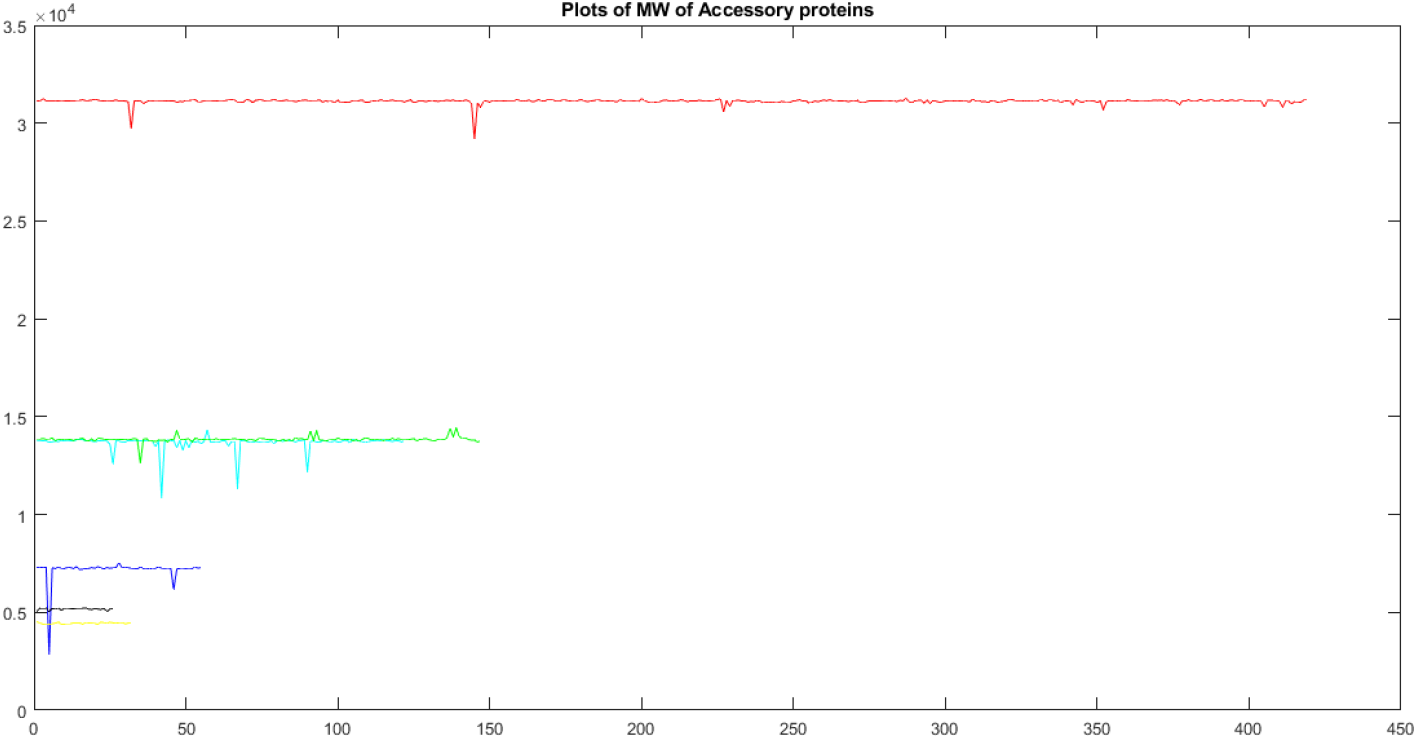
Graphical representations of molecular weights of unique accessory proteins

**Figure 13:**
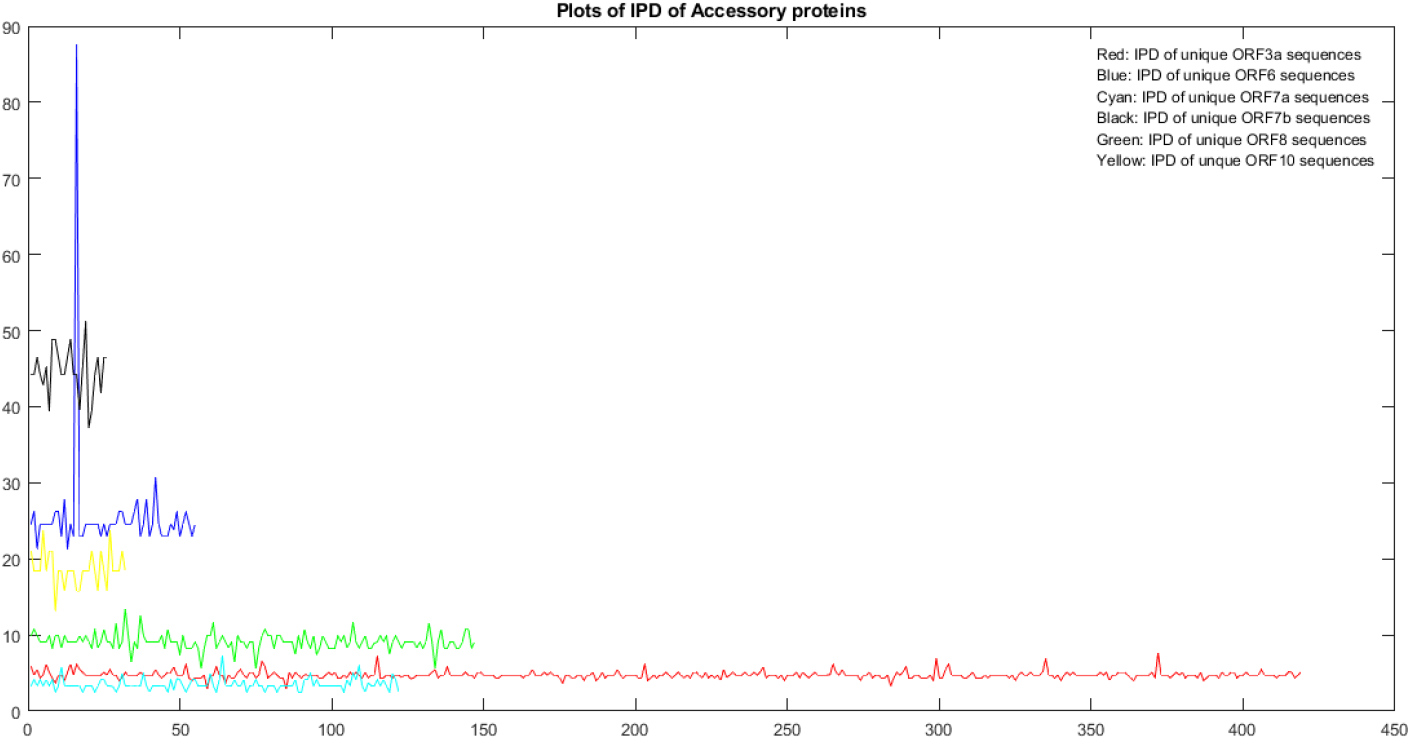
Graphical representations of Intrinsic Disorder contents of unique accessory proteins

**Figure 14:**
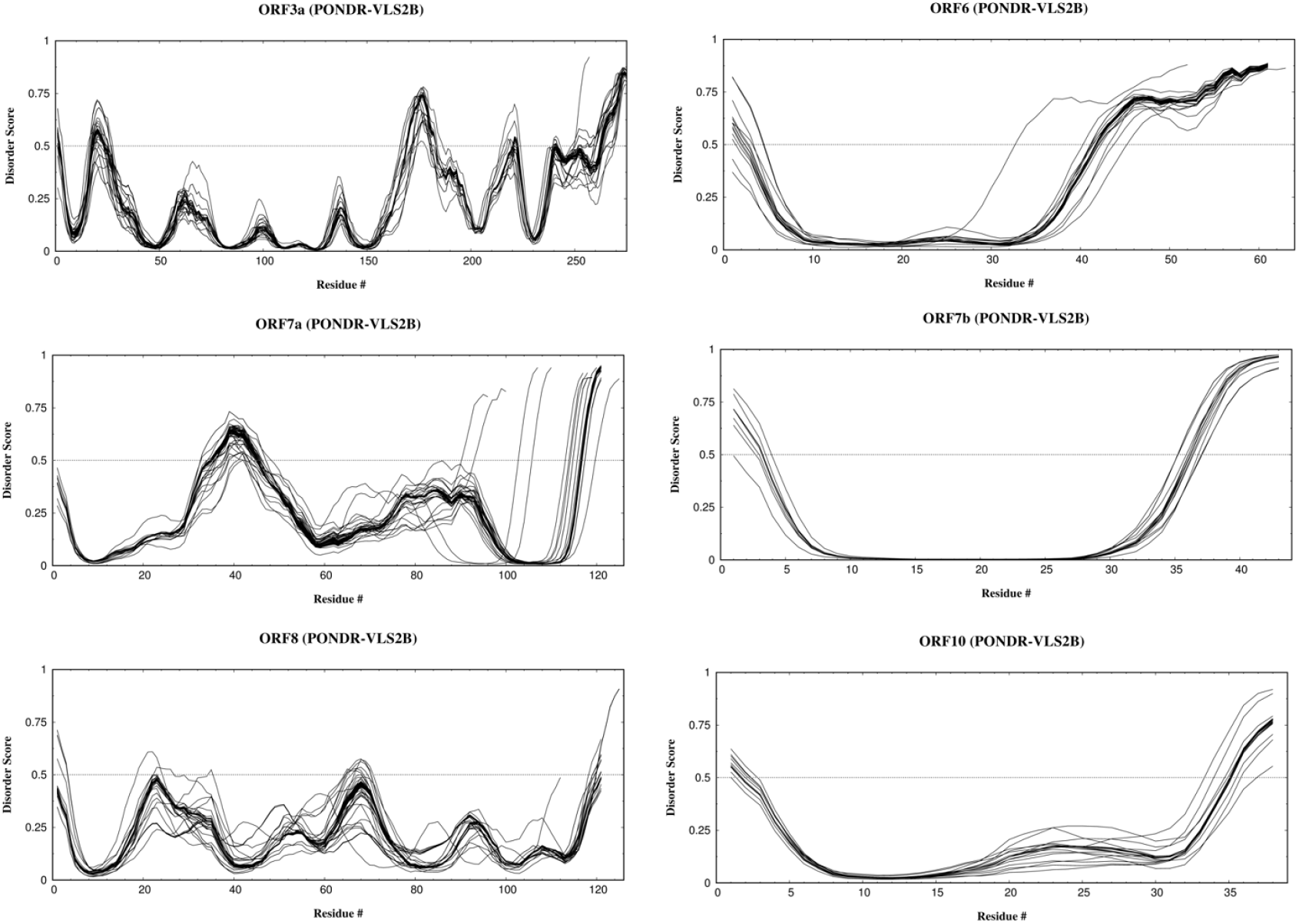
disorder plots of all unique variants for six accessory proteins of SARS-CoV-2

**Figure 15:**
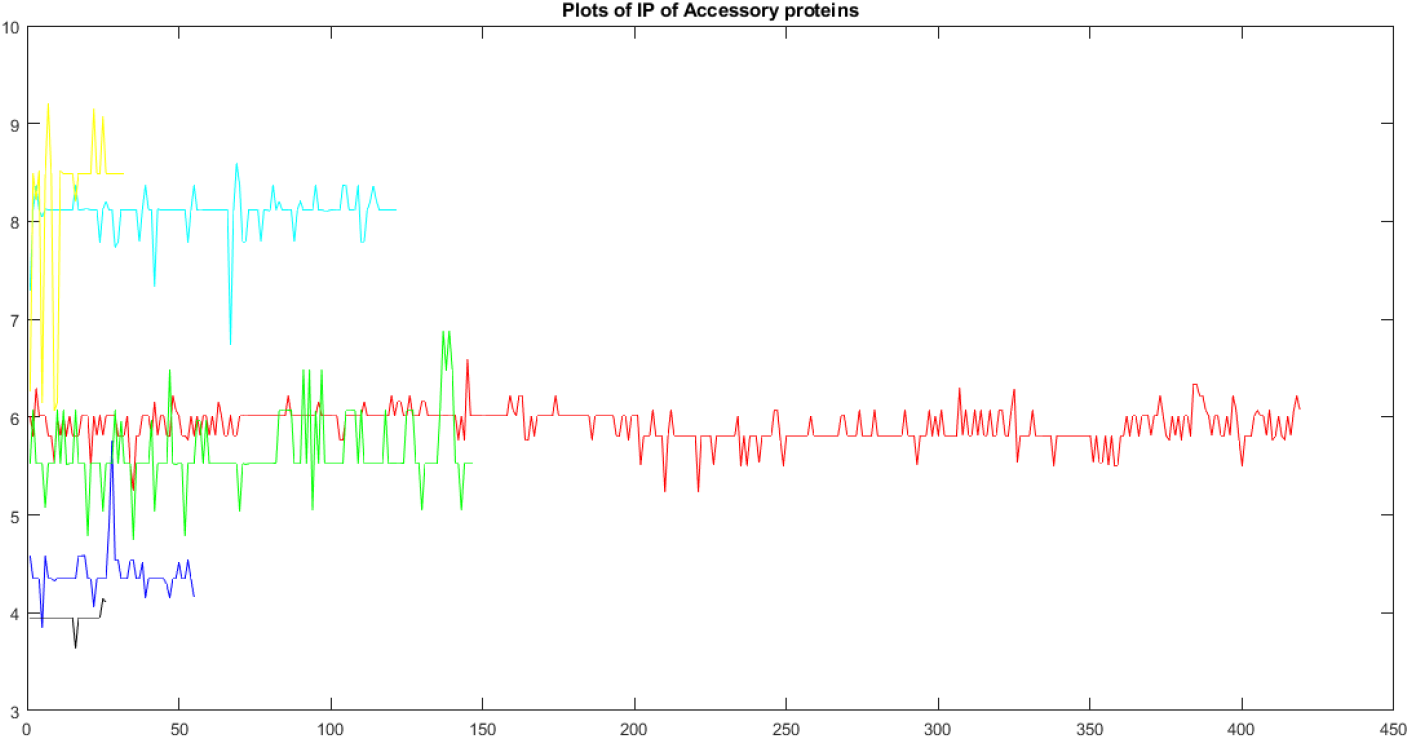
Graphical representations of isoelectric points of unique accessory proteins

## 4. Discussion

SARS-CoV-2 is the first HuCoVs with pandemic capacity due to its highly contagious nature deriving from the structural differences in its spike protein such as flat sialic acid binding domain, tight binding to its entry ACE2 receptor and capacity to be cleaved by furin protease [31]. However, 215 based on the estimated infection number close to one billion by WHO, SARSCoV-2 highly contagious but relatively a weak viral pathogen considering the overall of infection number has severe infections associated with the multiple organ dysfunctions [6]. This relatively weak pathological feature of SARS-CoV-2 could be related to the accessory proteins modulating host immunity as described above to the accessory proteins modulating host immunity as described above.

Based on the dynamic and various mutations on accessory protein variants, SARS-CoV-2 after diverging with BatCoV Ratg13 19 to 89 years ago was likely to have very few infections or somehow had very few selective pressure to tackle host immunity in nature [2]. In SARS-CoV-2 as other CoVs the genomic stability of their relatively large RNA genome around 30,000 amino acids is protected with proofreading proteins majorly 3’-5’ exonuclease non-structural protein (nsp)14 and 205 cofactor proteins nsp10, nsp13, and nsp16 [32]. Muller’s ratchet or also called as rachet effect explains the extinctive effect of high mutation rates of asexual organisms such as viruses [33]. Therefore, SARS-CoV-2 is repairing its mutations to preserve its genomic stability since a mutation can lead to pathological fitness losses or viral extinction [33]. However, there is a balance governed by genomic repair mechanisms such as nsp14 and viruses that require a certain degree of mutations to gain novel traits such as emergence transmission in zoonotic hosts [33]. For instance, SARS-CoV mutations, a 29-nucleotide deletion of ORF8, was associated with the less pathogenic strain [33]. Similarly, SARS-CoV-2 variants with a 382-nucleotide deletion, ORF8 had mild symptoms and did not require supplemental oxygen [33].

Furthermore, only one variant (identical to the Wuhan sequence (NC 045512) of each of the accessory proteins of ORF6, ORF7a, ORF7b, and ORF10 were present on all continents. Furthermore, it was observed that only two variants of ORF3a differed by a single mutation (Q57H, a clade/strain determining [30]) were found on all six continents. Also, in ORF8, only two unique variants (differed by a strain determining single mutation L84S) appeared on all continents. So, the maximally intersecting family of variations across all accessory proteins has turned out to be the smallest. These findings confirmed that the other variants of all the accessory proteins were due to demographic, environmental constraints. It was found that most of the unique variants of accessory proteins differed from the respective Wuhan accessory proteins by a single mutation, although basic descriptive statistics as found in section 3.1, unfolded their respective wide variability. Note that new variants of each accessory protein have been found in recent days and continue to do so. Significant amounts of unique variants of each accessory protein having wide variability might contribute significantly to the pathogenicity of SARS-CoV-2.

Therefore, our firm conviction that natural weakened stability (if achievable) of SARS-CoV-2 seems to be a far reachable destiny that alarms the danger of the present pandemic scenario due to COVID-19. Also, unique accessory protein variants across individual continents would all be expected to be mixed, while international travels would be restarted without strict protective measures. In this regard, it is our (SACRED, Self-Assembled COVID-19 Research & Education Directive”, consisting of international experts in mathematics, physics, computer science, bioinformatics, nanotechnology, structural biology, molecular biology, immunology, and virology) strong recommendation to governmental and non-governmental administrations to take necessary measures to mitigate the spread of COVID-19.

## Author Contributions

SSH conceived the project and carried out the preliminary work. SSH analyzed the results and wrote the primary draft of the article. All authors critically reviewed, edited, and approved the final manuscript.

## Supplementary Data

**Table 7:**
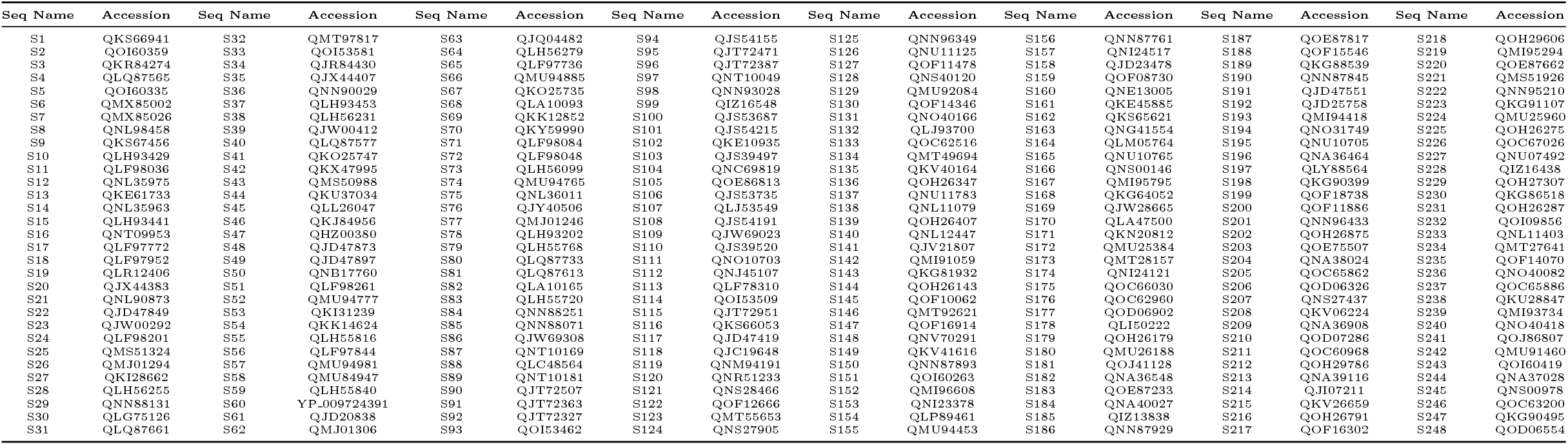
List of ORF3a Sequences and their corresponding accessions

**Table 8:**
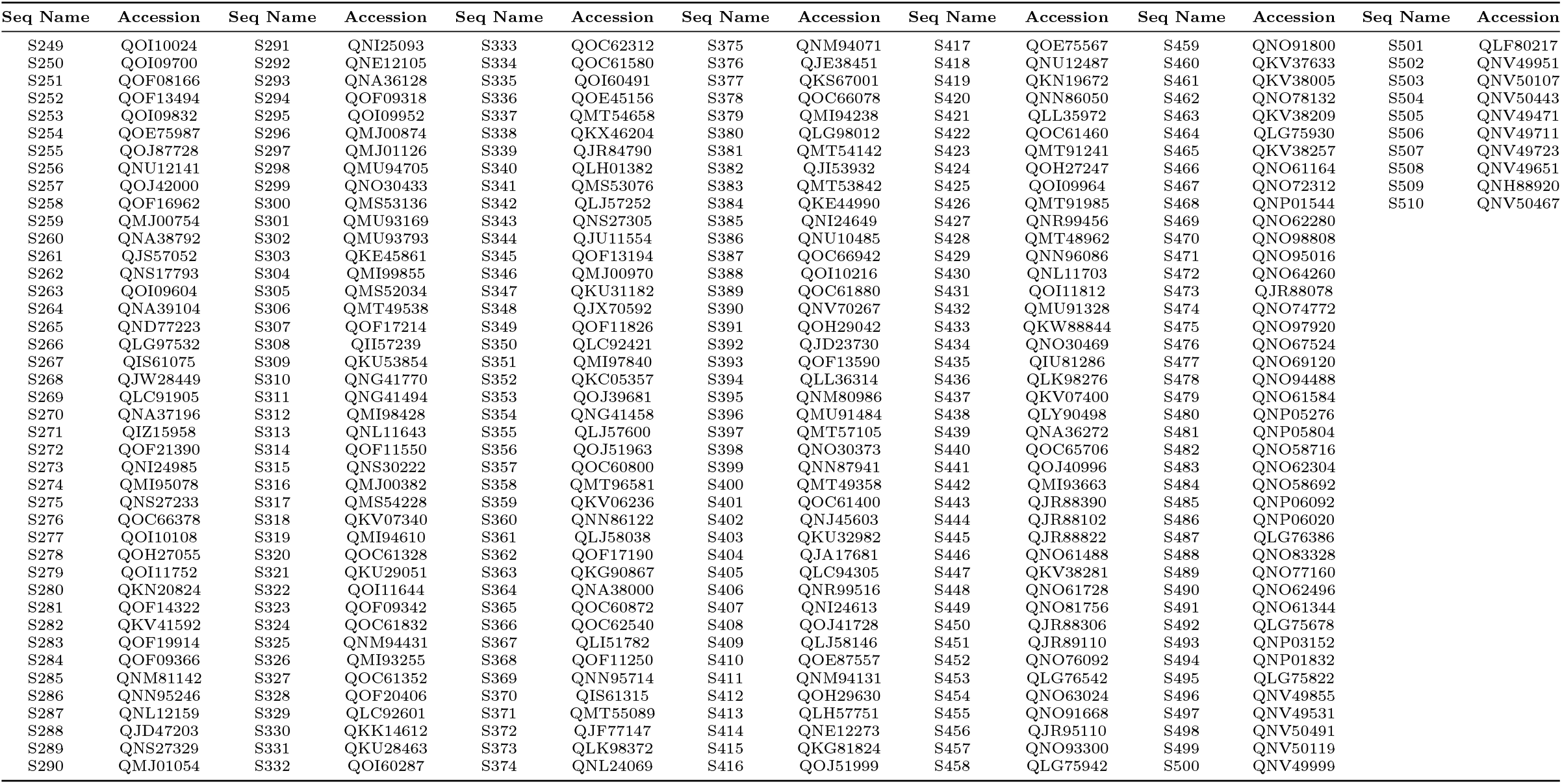
Contd … List of ORF3a Sequences and their corresponding accessions

**Table 9:**
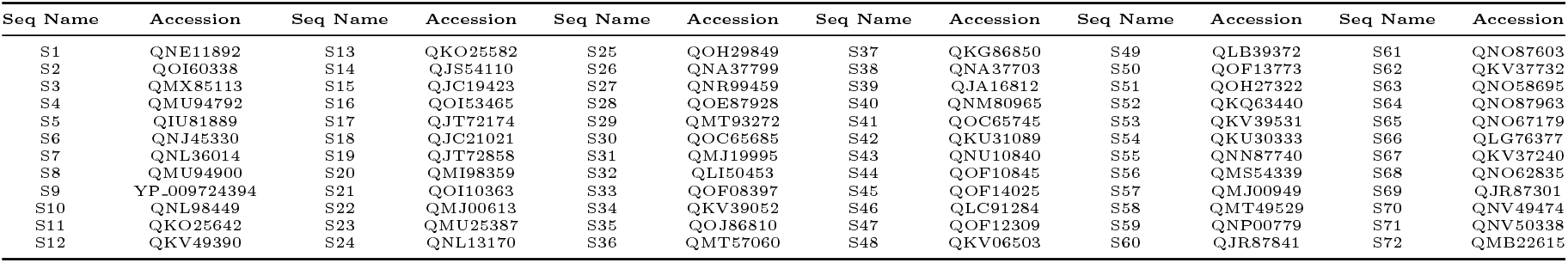
List of ORF6 sequences and their accessions

**Table 10:**
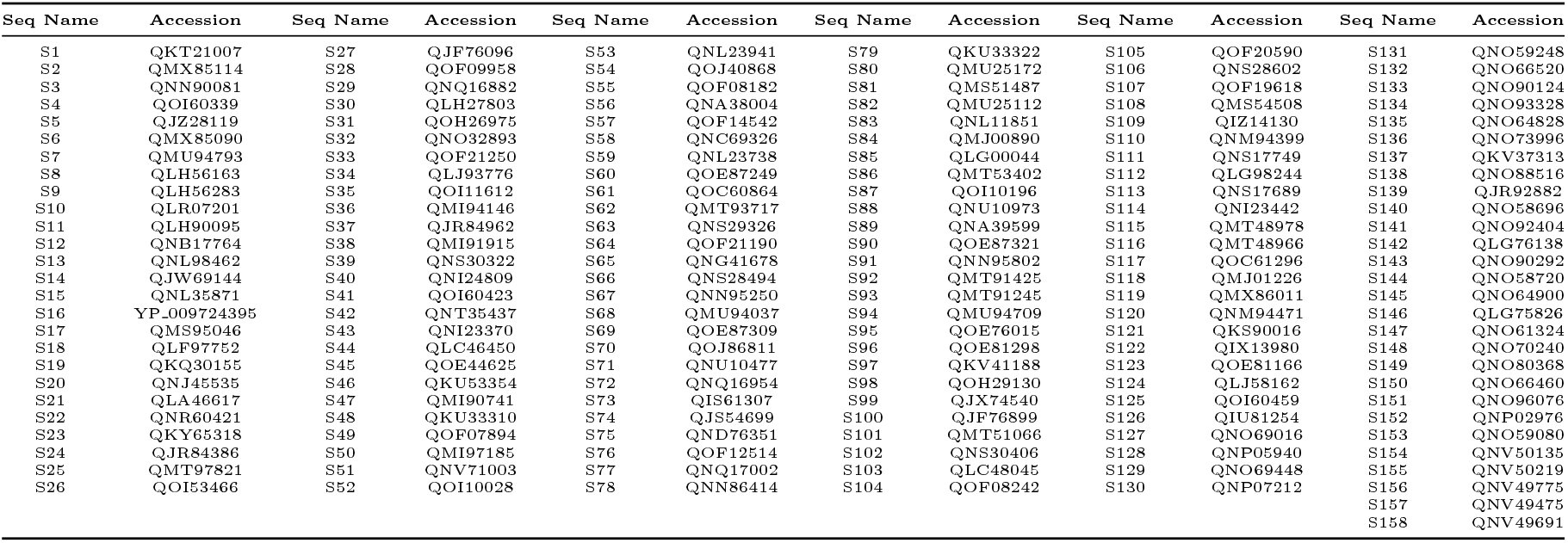
List of ORF7a sequences and their accessions

**Table 11:** List of ORF7b sequences and their accessions

| Seq Name | Accession | Seq Name | Accession | Seq Name | Accession | Seq Name | Accession |
| --- | --- | --- | --- | --- | --- | --- | --- |
| S1 | QNN90094 | S11 | QOI53467 | S21 | QOJ86812 | S31 | QNP01153 |
| S2 | QOI60340 | S12 | QNU10478 | S22 | QMI93943 | S32 | QLG75935 |
| S3 | QKX48960 | S13 | QNL24002 | S23 | QOH26412 | S33 | QNO58697 |
| S4 | QNJ45416 | S14 | QNL11060 | S24 | QNV70236 | S34 | QNO90425 |
| S5 | QMU84916 | S15 | QNU10274 | S25 | QNO30378 | S35 | QLG75923 |
| S6 | YP_009725318 | S16 | QMT53511 | S26 | QKE45866 | S36 | QNO74885 |
| S7 | QNN88304 | S17 | QOI10365 | S27 | QJC19833 | S37 | QNV49476 |
| S8 | QJQ84777 | S18 | QOF08027 | S28 | QKU28444 |  |  |
| S9 | QNL90926 | S19 | QMI96613 | S29 | QJD47604 |  |  |
| S10 | QKM76816 | S20 | QKY77886 | S30 | QKV38827 |  |  |

**Table 12:**
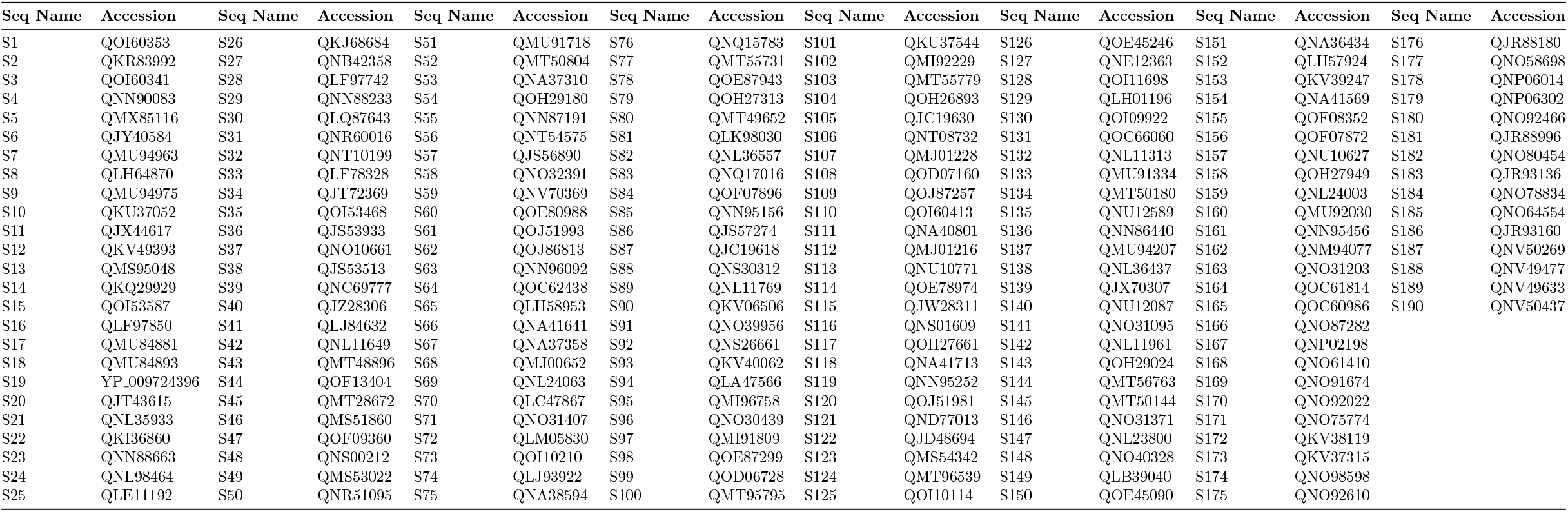
List of ORF8 Sequences and their corresponding accessions

**Table 13:**
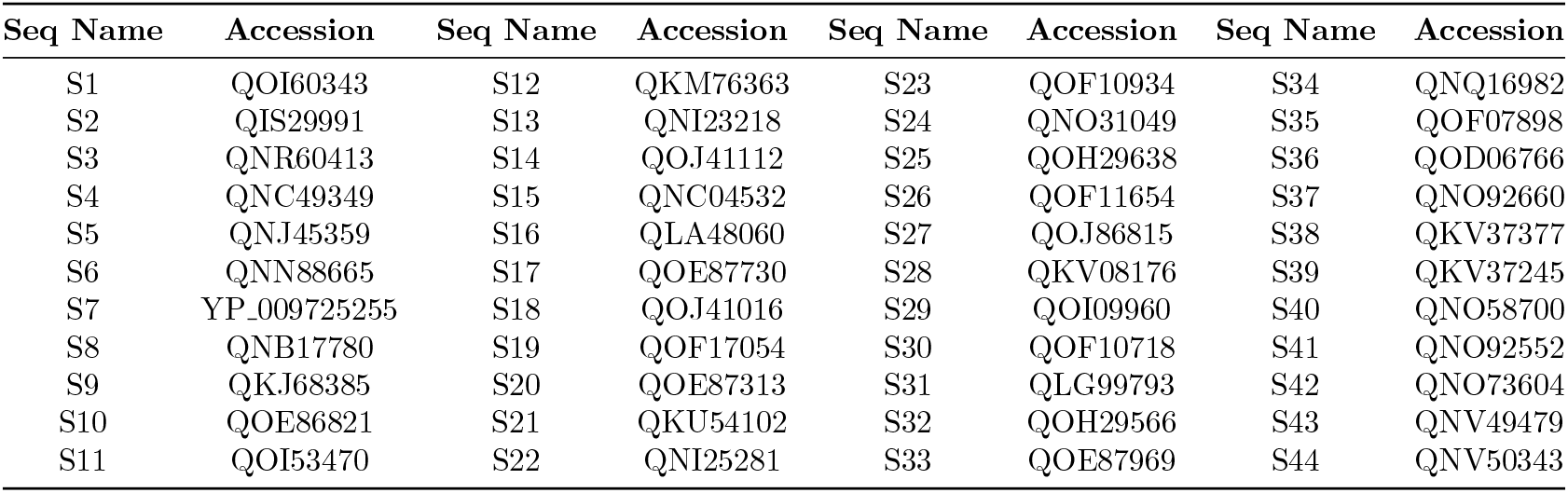
List of ORF10 Sequences and their corresponding accessions

